# CyTOF analysis identifies unusual immune cells in urine of BCG-treated bladder cancer patients

**DOI:** 10.1101/2020.06.12.147710

**Authors:** Eva Castellano, Célia Samba, Gloria Esteso, Laura Simpson, Elena Vendrame, Eva M. García-Cuesta, Sheila López-Cobo, Mario Álvarez-Maestro, Ana Linares, Asier Leibar, Thanmayi Ranganath, Hugh T. Reyburn, Luis Martínez-Piñeiro, Catherine Blish, Mar Valés-Gómez

## Abstract

High grade non-muscle-invasive bladder tumours are treated with transurethral resection followed by recurrent intravesical instillations of Bacillus Calmette Guérin (BCG). Although bladder cancer patients respond well to BCG, important questions remain unanswered, including how to identify at early stages non-responder patients and patients at risk to abandon the treatment. Here, we analysed the cells released into the urine of bladder cancer patients longitudinally 3-7 days after BCG instillations. Mass cytometry (CyTOF) analyses revealed that most cells were granulocytes and monocytes rather than effector lymphocytes, and most expressed activation markers. A novel population of CD15^+^CD66b^+^CD14^+^ CD16^+^ cells was very abundant in several samples and expression of these markers was confirmed using flow cytometry and qPCR. Samples of patients with a stronger inflammatory response contained more cells in urine; however, this was not due to haematuria, as the proportions of the cell populations observed were different from blood. We provide the proof-of-concept for a new approach to analyse samples that may help classify patients and identify those at risk of BCG infection and other unwanted BCG-related events.

## INTRODUCTION

Bladder cancer is the ninth most commonly occurring cancer in the world (Antoni *et al*, 2017) and the fifth most commonly diagnosed in Europe according to GLOBOCAN 2018 [Global Cancer Observatory (GCO), http://gco.iarc.fr/]. 70% of cases correspond to non-muscle invasive bladder cancer (NMIBC), that can recur and eventually become more aggressive and invade deeper layers of the bladder wall. Highgrade NMIBC (classified as Tis or Ta-1G3 tumours) is treated with transurethral resection followed by intravesical instillations of Bacille Calmette-Guérin (BCG), an attenuated strain of *Mycobacterium bovis* originally developed as vaccine for tuberculosis (Babjuk *et al*, 2017; Morales *et al*, 1976). The treatment was approved by the FDA decades ago and consists of weekly BCG instillations into the patient’s bladder, so that mycobacteria are applied directly to the tumour site. Therapy duration varies between different hospitals, but always comprises an induction phase of 6 weekly instillations and a maintenance phase comprising at least one year of 2-3 weekly instillations alternating at 3, 6 and 12 months. BCG represents the first successful immunotherapy, since the majority of patients respond well to this treatment and are free from relapse for at least three years after treatment. However, 30 to 40% of patients experience recurrence or progression and there are still no predictive methods for an early identification of nonresponder NMIBC patients (Gandhi *et al*, 2013; Kamat *et al*, 2017; Sylvester, 2011). Usually, failure of treatment is identified when the tumour has grown enough to either generate a positive urine cytology or a macroscopic mass that can be seen by direct inspection of the bladder with cystoscopy or by means of imaging with ultrasound or CT scan. Thus, the identification of biomarkers that allow early identification of non-responder NMIBC patients is a clear unmet need. Additionally, although BCG treatment is effective, a high number of patients suffer adverse effects due to the presence of living bacteria in the bladder, such as irritation or mild fever, and a small percentage of them can even suffer systemic problems associated with the treatment (Lamm *et al*, 1992; Perez-Jacoiste Asin *et al*, 2014), such as sepsis, pulmonary or urogenital infections. Research is still needed in this area to prevent these severe side effects and to adjust the adequate BCG dose to each particular patient depending on their immune response (Larsen *et al*, 2019).

Several clinical and pathological parameters as well as soluble immune factors have been proposed to have value as prognostic markers for NMIBC patients. Urologists usually classify patients following models widely known for prediction of tumour recurrence and progression, and risk tables have been elaborated by the European Organization for Research and Treatment of Cancer (EORTC) (Cambier *et al*, 2016; Sylvester *et al*, 2006) and the Club Urológico Español de Tratamiento Oncológico (CUETO) (Fernandez-Gomez *et al*, 2009). These models are based on clinical parameters such as histological grading, pathological stage, tumour multiplicity, etc, but their predictive power is still insufficient.

Laboratory-based research has proposed using other variables such as the amount of cytokines in urine (Ashiru *et al*, 2019; Bisiaux *et al*, 2009) or the combination of the concentration of soluble factors in urine with clinical data (CyPRIT) (Kamat *et al*, 2016). However, their use is still not in place in the clinics since studies in larger cohorts are needed.

Due to difficulties in studying immune responses to tumours in humans, many laboratories have addressed these questions in animal models or *in vitro* experiments. In these studies, researchers have concluded that instillations with BCG result in immune cell infiltration in the bladder as a consequence of the inflammatory response (Pettenati & Ingersoll, 2018; Redelman-Sidi *et al*, 2014). *In vitro* studies have investigated the effect of BCG on different immune cell types; granulocytes (Suttmann *et al*, 2006), dendritic cells (Kumar *et al*, 2018) and natural killer (NK) cells (Brandau & Bohle, 2001; Brandau *et al*, 2001; Garcia-Cuesta *et al*, 2015), which become activated in response to mycobacteria. For example, we have recently described an anti-tumour specific population of CD56^bright^ CD16^+^ NK cells after *in vitro* exposure of peripheral blood mononuclear cells (PBMCs) to BCG that display cytotoxic activity against bladder cancer cells (Garcia-Cuesta *et al*, 2017). Pioneering studies 30 years ago revealed that the number of cells in urine significantly increased 24 hours after BCG administration and that neutrophils were a predominant cell type (De Boer *et al*, 1991). However, no data is available on which cells are released into the urine after the initial phases of cystitis and haematuria have resolved. We hypothesised that healing immune cells are recruited to the tumour site several days after BCG instillations and their analysis could provide a new tool to follow the intensity of the immune response in bladder cancer patients. This idea is further supported by our recent finding of cytokines, in particular CXCL10, released into urine seven days after BCG instillations, in higher amounts in patients with more side effects (Ashiru *et al*., 2019).

To directly study the immune response occurring in bladder cancer treated with BCG, we planned a detailed analysis of cells recruited to the bladder wall and secreted in the patient’s urine in a longitudinal study obtaining samples at multiple time points after BCG instillations. Besides the fact that urine is an ideal non-invasive source of biomarkers, in the case of bladder cancer it provides information on changes in the tumour environment. To this end, we established and optimised several methods, including traditional flow cytometry and mass cytometry (CyTOF). Given that immune cells are absent from the urine of healthy donors, it is not possible to directly compare urinary cells of BGC-treated patients with healthy controls. However, it was possible to analyse cells that appeared in patients in response to the BCG treatment and these cells theoretically would have been in close contact with the tumour. Microscopy revealed both immune- and epithelial-shaped cells in most patients. Flow cytometry optimization was challenging, due to hypoxia and autofluorescence. However, after overcoming these problems, several panels for staining of urinary cells could be established. CyTOF revealed that urinary cells were primarily comprised of myeloid and granulocytic lineages. Interestingly, most cells expressed markers of activation, such as CD69 and CD107a. Some samples also contained effector lymphocytes, T cells and NK cells, although the cell numbers of these populations were generally low. A CD15^+^CD66b^+^CD14^+^ granulocyte population was identified in several samples from patients with extremely elevated cell infiltration. These cells resemble APC-like tumour associated neutrophils (TAN) or myeloid-derived suppressor cells (MDSC). Gene expression of population markers was also validated by quantitative PCR (qPCR), in particular the granulocyte marker CD66b, the myeloid marker CD14, as well as the NK marker CD56.

In summary, this study is a proof-of-concept that a combination of clinically feasible techniques such as flow cytometry and qPCR can be used for systematic studies of the immune response elicited after treatment of NMIBC patients using BCG. These parameters, in combination with cytokines and clinico-pathological data, may help to establish the different levels of response and so aid urologists to classify patients as responder, non-responder or prone to adverse events. This, together with further studies will allow the combination of all these data to be used for early classification of patients.

## RESULTS

### Bladder cancer patient urine contains immune cells after BCG instillation

Bladder cancer patients start BCG instillations a few weeks after transurethral resection, if haematuria has resolved. In a previous study of cytokines released to urine during BCG treatment (Ashiru *et al*., 2019), a clear increase in urine cells was observed once instillations started. Although it was possible to count a few thousand cells per ml of urine in some patients at the beginning of treatment, most patients had urinary cell numbers above 10^4^/ml only after they had started receiving BCG instillations. A new study was designed to explore whether the identity of the cells recruited to urine could provide information on the efficacy of the immune response stimulated by mycobacteria. Because the treatment involves many cycles of instillations, when possible, several samples were obtained from the same patient after different cycles of instillations. Because we were interested in the cells recruited after the initial inflammation has been resolved, samples were obtained between 2 and 7 days after instillations (Supplementary Table 1). The number and shape of urinary cells varied between donors, however, hematopoietic-like, as well as flat, fibroblast-like cells could be observed consistently by light and electron microscopy in these preparations (Figure 1A).

**Figure 1.**
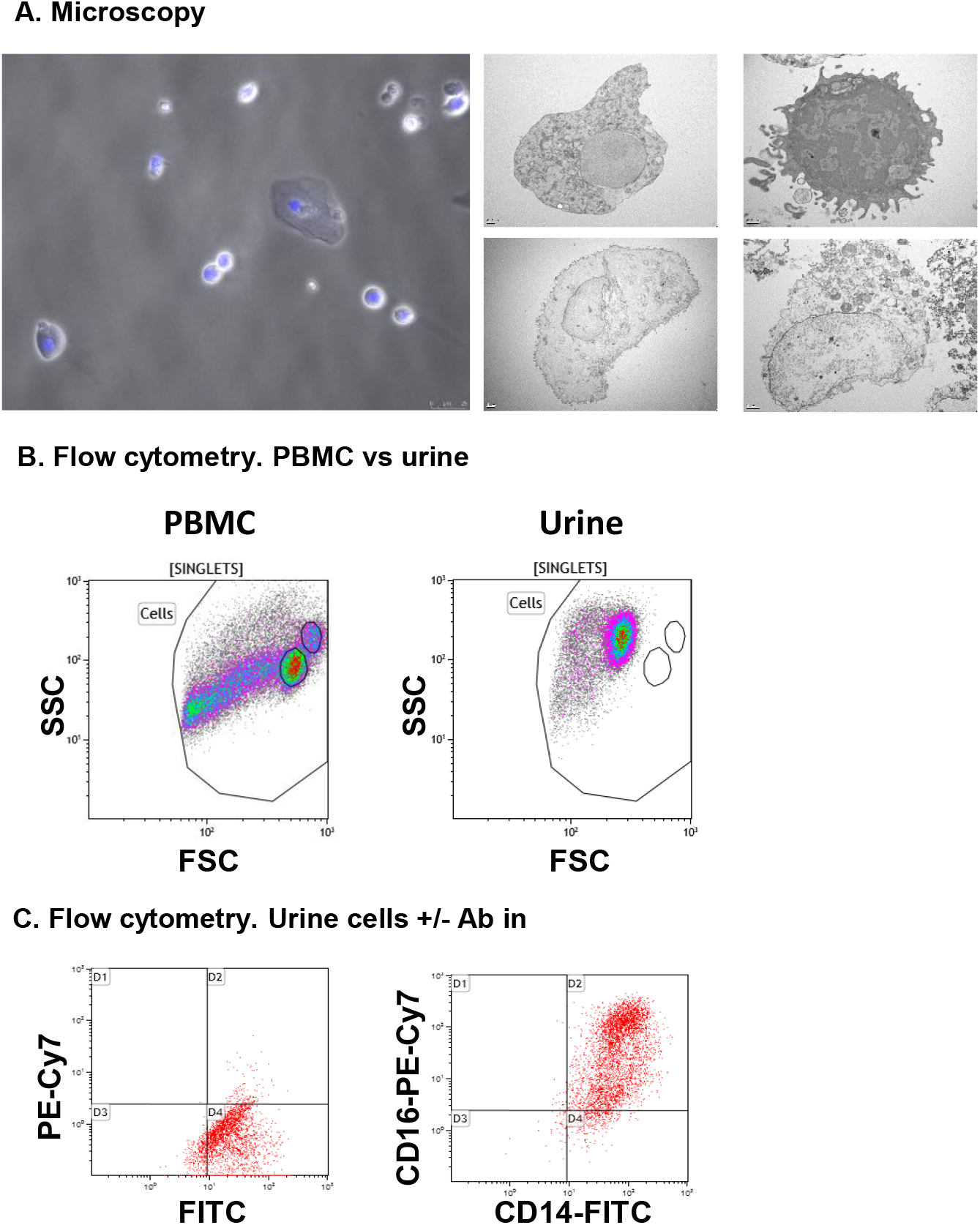
Cells recruited to urine analysed by microscopy and flow cytometry. Cells from urine of bladder cancer patients receiving intravesical instillations of BCG (Ta/T1G3 or CIS, mean age 68.5 years) were analysed. **A. Microscopy.** For optical microscopy (left), cells were immobilised on cover-slips using poly-L-lysine and nuclei were stained using DAPI. Samples were visualised in a fluorescence microscope. For electron microscopy (right panels), cells were immobilised in nickel grids and sections stained with uranyl acetate and lead citrate. The pictures show examples of cells with different morphology and integrity observed in different patients. **B. Comparison of size and complexity by flow cytometry of peripheral blood mononuclear cells (PBMC) from buffy coats and cell recruited to urine.** PBMC from healthy donors were isolated and analysed by flow cytometry. Plots represent the Forward Scatter (FSC) vs Side Scatter plots (SSC) keeping the region gates in the same position for both PBMCs from buffy coats and cells recruited to urine. **C. Autofluorescence in urine recruited cells.** Cells were either kept unstained (left) or stained (right) with the indicated antibodies and cell surface staining was analysed in comparison.

To investigate the nature of the cellular immune response occurring in BCG-treated NIMBC patients, initial experiments were designed to assess the phenotype of urinary cells by multiparametric flow cytometry. Samples were collected several days after instillation with BCG, thus allowing the evaluation of cell populations persisting after the initial inflammation. Initial analyses of Forward Scatter/Side Scatter (FSC/SSC) plots showed that urinary cells differed from those in peripheral blood, presumably due to the different osmolarity and pH conditions (Figure 1B). Despite the unsightly flow cytometry plots, it was clear that certain antibodies were staining the cells; in particular, CD14, and CD16 were easily observed in many of the samples analysed (Figure 1C). These results suggested that urine immune cells could be a useful source of information about the level of inflammation and anti-tumour response of bladder cancer patients several days after treatment with BCG. Thus, it was important to optimise a method to evaluate multiple parameters in these cell populations.

Flow cytometry experiments posed a number of technical problems, which could be due to the hypoxic conditions of urine to which the cells had been exposed for several hours prior to staining. Some antibodies failed to stain and different clones were carefully selected. Additionally, the events acquired in the flow cytometer appeared to have a small size and autofluorescence was very high. To address these issues, several experiments were carried out to optimise the methodology for flow cytometry of cells in urine. In hypoxic environments, like urine, cells have their metabolism skewed towards glycolysis and accumulate nicotinamide dinucleotides (NADH and FAD) which have fluorescence peaks in the range of the fluorophores used for flow cytometry (NADH Ex Max = 350nm; FAD Ex Max = 450nm; NADH Em Max = 450nm, FAD Em Max 530nm) (Palmer *et al*, 2015). Thus, fluorescence artefacts were reduced by turning off the green channels in the flow cytometer, so that endogenous fluorescence would not interfere. In parallel, DNA content was determined by staining with Hoechst dye and a gating strategy was established using all this information (Supplementary Figure 1A). To validate the specificity of antibody staining, blocking experiments were carried out. Non-specific staining was not observed, since blockade of Fc receptors with either human Ig or human serum did not decrease signal (Supplementary Figure 1B).

These experiments demonstrated that urine samples from bladder cancer patients contained immune cells that could be further characterised by flow cytometry.

### Mass cytometry can be used to detect and phenotype urinary cells in BCG-treated patients

In order to explore a large number of markers using the limited number of cells available per patient, urinary cells were analysed by CyTOF, a technique that allows the detection of 40+ markers simultaneously at the single cell level. An additional advantage of CyTOF is that it does not rely on fluorescence, and thus allowed us to overcome the technical challenges caused by the autofluorescence observed in the conventional flow cytometry experiment.

CyTOF was used to perform a comprehensive study of multiple parameters in urinary cells from different bladder cancer patients at different time points. Due to limitations in sample acquisition rate and the expense associated with CyTOF, only the initial cycle of instillations from a limited number of patients was analysed and used as a tool to inform further flow cytometry studies.

Since there were no reports on handling urinary cells for mass cytometry analysis, the first challenge was to confirm whether these hypoxic, previously frozen urine cells could be analysed using this technology. To do so, two samples from different donors were thawed into cold RPMI containing 20% FCS and benzonase. Cell viability was above 80% and aliquots of 0.5-2·10^6^ urine cells were stained for mass cytometry using a panel of 35 antibodies. Because our preliminary flow cytometry data revealed a large number of CD15^+^ and CD16^+^ cells, a panel of antibodies designed to detect mainly myeloid cells markers as well as markers for lymphocyte lineages was used (Supplementary Table 2). Plots from urine were compared to PBMC and to a mixture of PBMC and purified neutrophils (not shown).

To assess cell composition by CyTOF, intact cells were gated based on ^191^Ir (DNA-1) vs ^193^Ir (DNA-2). This was followed by a live/dead stain based on cisplatin (Fienberg *et al*, 2012). An example of the gating strategy is shown in Figure 2A. Since the membrane integrity of urinary cells was expected to be affected by urine, a less stringent gating for the live/dead stain was used for urine cells. The identification of immune populations was performed manually by comparing blood samples with urinary cells (not shown). These comparisons indicated that we could identify the major cell lineages by CyTOF in urinary cells.

**Figure 2.**
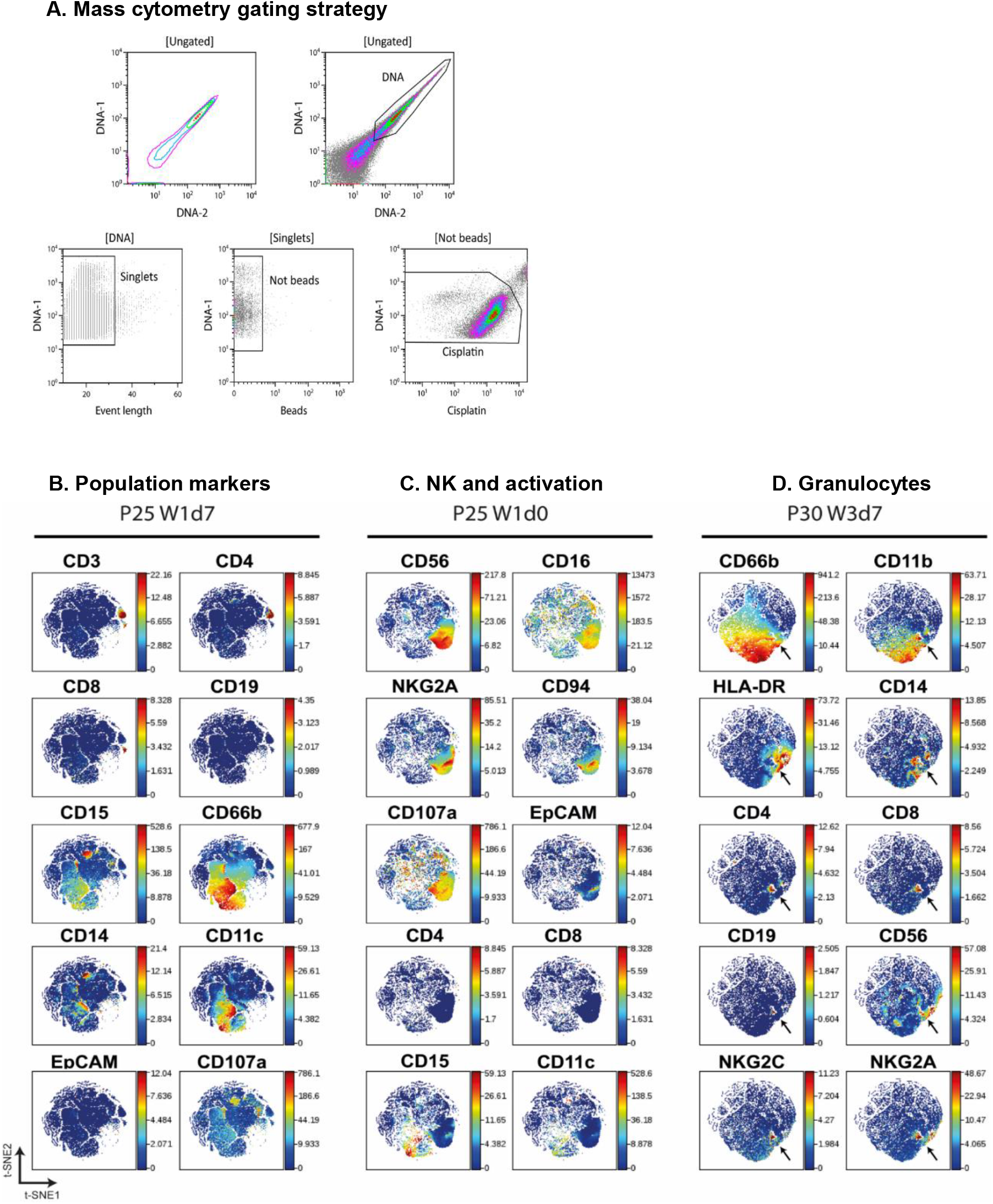
Mass cytometry on urine cells. **A. Mass cytometry gating strategy**. A first region on the DNA positive events was selected to gate singlets. Calibration beads and cisplatin positive events were then sequentially excluded. **B. C. D. Validation of antibody staining by mass cytometry.** viSNE analysis were performed separately for different individuals (patients 25 and 30) and several plots were compared to visualise the markers for different immune populations. Arrows indicate an example of a population binding antibody non-specifically.

Once it had been established that CyTOF could be used to phenotype cells released in the urine of BCG-treated NMIBC patients, 44 samples obtained at different time points of the treatment from 12 different BCG-treated bladder cancer patients were stained and analysed by CyTOF (Table 1). The data obtained in different days were normalized (Supplementary Figure 2), and viSNE was used to visualize the distribution of cell phenotypes (Amir el *et al*, 2013)(See methods). Several viSNE analyses were done, grouping different samples in each case, so that we could compare initially the samples corresponding to a given treatment day, and then grouping the samples corresponding to a particular donor.

**Table 1.**
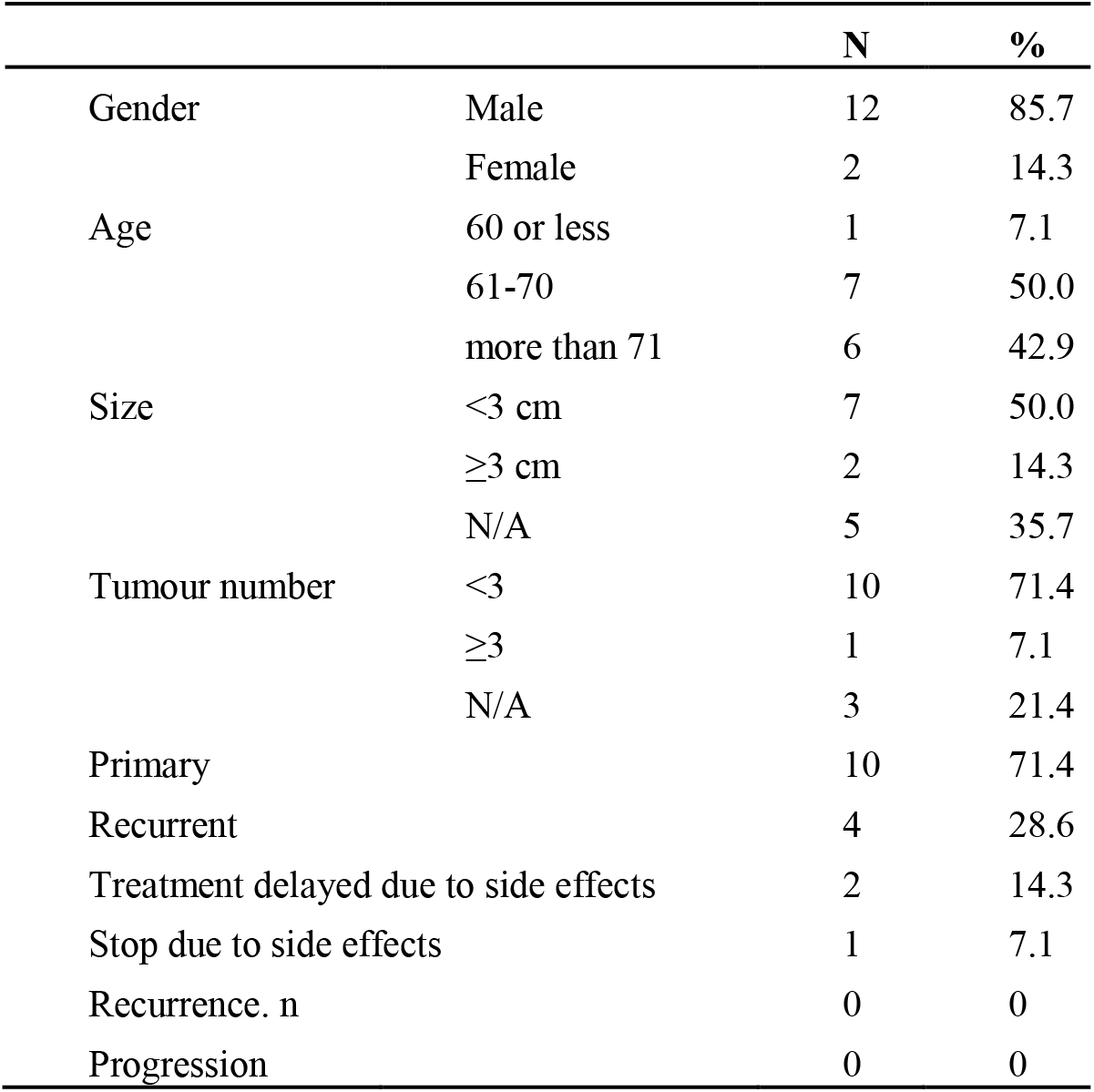
Patient demographics and clinical features.

|  |  | N | % |
| --- | --- | --- | --- |
| Gender | Male | 12 | 85.7 |
|  | Female | 2 | 14.3 |
| Age | 60 or less | 1 | 7.1 |
|  | 61-70 | 7 | 50.0 |
|  | more than 71 | 6 | 42.9 |
| Size | <3 cm | 7 | 50.0 |
|  | ≥3 cm | 2 | 14.3 |
|  | N/A | 5 | 35.7 |
| Tumour number | <3 | 10 | 71.4 |
|  | ≥3 | 1 | 7.1 |
|  | N/A | 3 | 21.4 |
| Primary |  | 10 | 71.4 |
| Recurrent |  | 4 | 28.6 |
| Treatment delayed due to side effects |  | 2 | 14.3 |
| Stop due to side effects |  | 1 | 7.1 |
| Recurrence. n |  | 0 | 0 |
| Progression |  | 0 | 0 |

Before examining viSNE data in detail, the maps were used to verify antibody binding specificity and to confirm that the markers expressed by different cell populations were in general consistent with the molecules expected to be expressed by particular lineages (Figure 2B-D). As expected, the CD66b population, corresponding to CD15^+^ granulocytes, did not express lymphocyte markers such as CD19, CD3 or CD56 or the myeloid marker CD14, that co-localised with CD11c. Similarly, CD3^+^ T lymphocytes could be divided into CD4^+^ and CD8^+^ and these populations were negative for monocyte or granulocyte markers. EpCAM was not expressed in hematopoietic cells. CD56^+^CD16^+^ NK cells also expressed the NK markers CD94, NKG2A, but not CD8 or CD4 (Figure 2C). In some cases, a small population that showed nonspecific binding of many different lineage markers was identified, suggesting an artefact of staining. These cells were eliminated from further analyses (Figure 2D).

Thus, although cells recruited to urine had suffered extreme conditions, their phenotype could still be analysed using CyTOF.

### Urine obtained from BCG-treated patients at weeks 0 and 6 contains heterogeneous cell populations

To have a first broad view of the different cell types that could be found at different stages of the BCG treatment, samples obtained before any instillation (time zero) (Figure 3A) and before the 6^th^ instillation (Figure 3B) were compared using viSNE. Most patients did not have cells before starting the instillations and only 3 samples corresponding to time zero were available to analyse cells (Supplementary Table 1).

**Figure 3.**
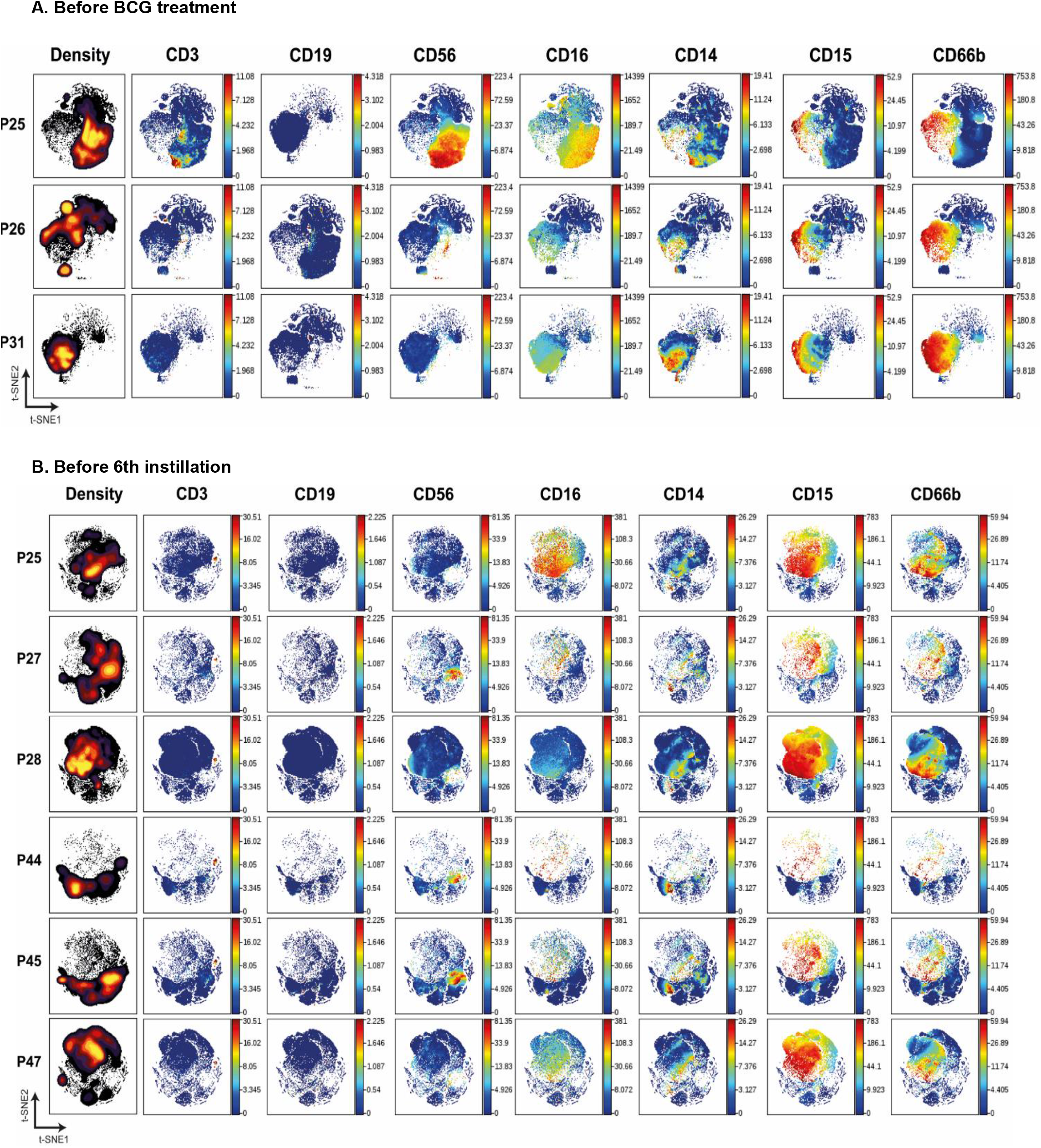
Mass cytometry analysis of cells from patients at the same stage of treatment. viSNE analysis was performed grouping patients at the same stage of treatment: either before starting BCG treatment (A) or 7 days after receiving the 5^th^ instillation (B). The left column represents the density plots while the rest of the plots correspond to the main PBMC population markers.

The presence of cells in urine at this stage is probably due to the inflammatory process and healing of the post-surgical bladder wound. It is unlikely that these cells correspond to simple haematuria because the percentages of leukocytes observed in these samples were not consistent with the composition of blood populations.

The abundance of different cell populations was represented in density maps and the relative amount of the different population and activation markers were visualised in colour maps (Figure 3). At time zero, the distribution of cells analysed in the three different patient samples differed. One patient (P25) had a considerable population of lymphocytes, mainly NK cells and some CD3^+^ lymphocytes, while the other two patients (P26 and P31) had primarily granulocytes and monocytes (Figure 3A).

Before receiving the 6^th^ instillation, myeloid and granulocyte populations predominated in all patients (Figure 3B). A small percentage of monocytes and even smaller proportions of CD3^+^ lymphocytes and NK cells could be detected in most samples. B cells were generally absent. Cells not binding any of the antibodies used against immune populations, probably epithelial cells, were also abundant and could be visualised by electron microscopy (Figure 1A).

### CD66b^+^ granulocytes in urine of bladder cancer patients are a heterogeneous population

The first analyses, comparing two different time points of the BCG treatment, revealed the general characteristics of the cells released into urine of bladder cancer patients, with some heterogeneity among samples. However, a second level of heterogeneity can be considered in this study, that is donor-to-donor variation. To further explore the combination of different markers and populations present in urine samples from one patient during different stages of the treatment, viSNE analyses were carried out after grouping all the available samples for each one of the patients (Figure 4A, top rows). This second analysis was done to evaluate the effect of time in cell heterogeneity for each donor.

**Figure 4.**
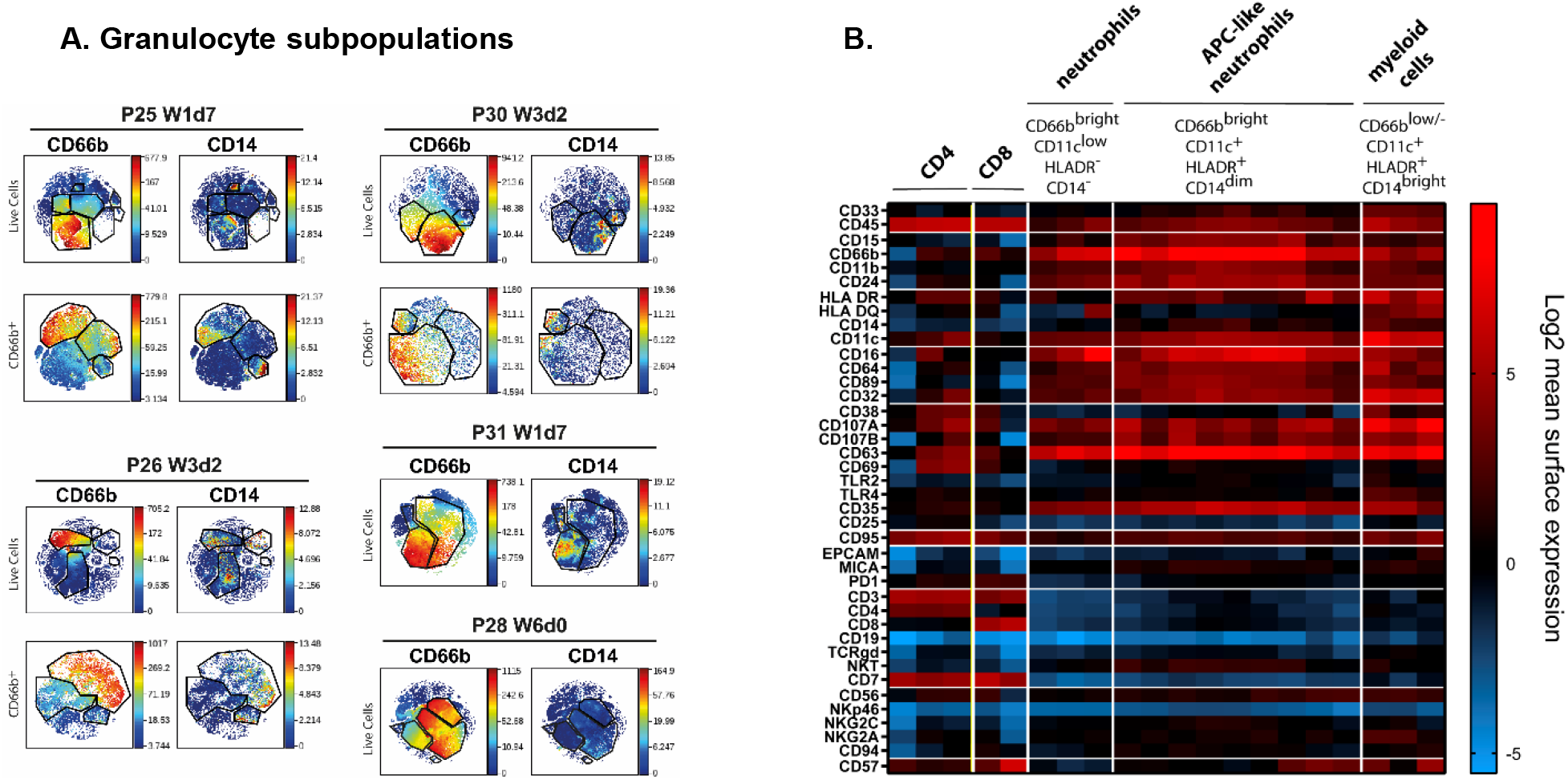
Mass cytometry analysis of cells from patients at the same stage of treatment. viSNE analysis was performed grouping all the samples analysed from each patient during the first cycles of BCG treatment. **A. viSNE analysis of granulocyte populations.** Granulocyte populations included several subpopulations that expressed distinct surface markers. For a better visualization of these subpopulation, viSNE was performed gating on CD66b (bottom rows). **B. Heat map.** To aid visualisation, the combinations of markers defining the major populations of leukocytes observed in urine of different patients were plotted. This plot allows to identify the different markers expressed in the different populations identified. The figure represents several samples obtained from different patients.

It was surprising to identify CD66b^+^ cells that co-expressed CD14 and CD11c in many different patients. Further, it was evident that the abundant granulocyte population was composed of several subpopulations which expressed different levels of other molecules such as CD16, HLA-DR and CD11c. For better resolution of this unexpected population, we first gated on CD66b^+^ cells and then ran viSNE analysis just on this population for each patient (Figure 4A, bottom rows). It was then possible to identify CD14^+^ cells within the gate of CD66b^+^ granulocytes.

To explore the different markers expressed by CD66b^+^ cells, a heat map was built and used to compare the markers expressed by CD66b^+^ and by lymphocyte populations (Figure 4B). This was done as an exercise to help classify cell populations and to identify similar populations in different samples and patients; thus, this heat map does not show a proportional clustering of the populations enriched during treatment, but rather is a tool to help visualising the patterns of expression contained in the large number of samples analysed. Using this visualization tool, it was confirmed that CD3^+^ cells also expressed CD45 and CD7, could express either CD4 or CD8, and were negative for other markers such as CD19, CD66b, CD11b or CD11c. Myeloid cells were CD14^+^ CD11c^+^ HLA-DR^+^, but CD15^-^ and CD3^-^, while CD66b^+^ cells could appear either as classical granulocytes CD14^-^ HLA-DR^-^ CD11c^lo^, or CD14^+^ HLA-DR^+^ CD11c^+^, all of them being negative for lymphocyte markers. It was also possible to identify populations with non-specific binding of antibodies that were excluded from any further analyses. Interestingly, CD66b^+^ CD15^+^ cells also expressed CD107a and CD69, corresponding to activated granulocytes. Different levels of CD66b expression resulted in two types of granulocytes that were designated *dim* and *bright;* however, they also could be distinguished by different levels of HLA-DR, CD16 and CD11c expression. Unexpectedly, a hybrid population with features of both granulocytes and monocytes (CD14^+^ CD15^+^ cells) was also identified in these samples. CD14^+^ CD15^+^ cells have been described previously in cancer patients. This phenotype could correspond to antigen presenting cells-like hybrid tumour associated neutrophils (APC-like TAN) in which markers for both granulocytes (CD11b, CD66b, CD15) and monocytes (CD33, CD14, HLA-DR, CCR7, CD86, CD20) co-exist.

T lymphocytes, expressing CD45, CD7, CD3 and either CD8 or CD4, were only found in a few samples. Similarly, only a few samples contained NK cells. When detected, lymphocytes were usually positive for activation markers and the combination of markers showed consistency within each population.

### APC-like tumor associated neutrophils were identified in freshly isolated urine cells

Since several populations observed in CyTOF data using viSNE revealed unexpected combinations of markers (Supplementary Figure 3A), these phenotypes were confirmed by manual analysis (Supplementary Figure 3B) and new flow cytometry analysis (Supplementary Figure 3C). Additionally, samples from three new different patients receiving the 6^th^ BCG instillation (Week 6), were also analysed by flow cytometry, revealing again three populations: CD15^+^ CD14^-^, CD15^+^ CD14^+^, CD15^-^ CD14^+^ (Figure 5). This new experiment, performed with freshly isolated urine cells from three patients in parallel allowed flow cytometry discrimination between CD14^+^CD15^+^ TAN and CD14^+^CD15^-^ monocytes. CD14^+^CD15^+^ TAN and CD14^+^CD15^-^ monocytes expressed CD11c and HLA-DR while the granulocyte population CD15^+^ CD14^-^ did not express these markers. Each subpopulation was also easy to identify in the FSC/SSC region and was present in different amounts in the three patient samples stained in parallel. These data are consistent with those obtained by CyTOF. Looking at the combinations of CD66b, CD16, CD11c and HLA-DR, several cell populations were identified: monocytes CD66^-^ CD11c^+^ HLA-DR^+^ CD16^lo^; granulocytes CD66b^bright^ CD11c^-^ HLA-DR^-^ CD16^+^; and APC-like neutrophils CD66b^bright^ CD14^dim^ CD11c^+^ HLA-DR^+^ CD16^+^. It was also clear by flow cytometry that different patients have different percentages of each population within total urine cells. The percentages of immune populations obtained were manually gated to confirm the subpopulations identified (Supplementary Figure 4). Interestingly, this analysis revealed that a large number of cells were not stained with the immune population markers, and these cells likely correspond to either tumour cells or epithelial cells from the genito-urinary tract.

**Figure 5.**
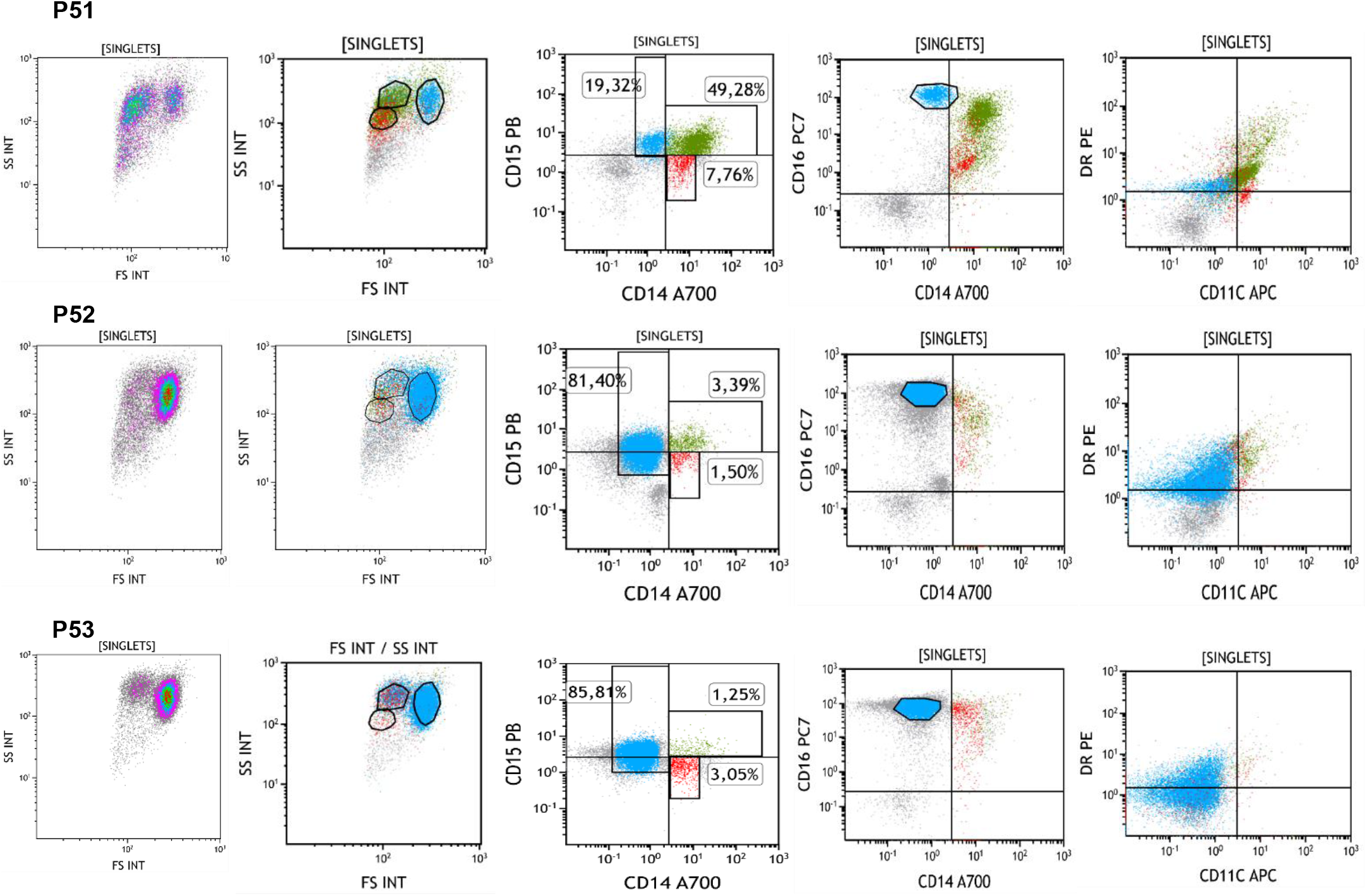
Different populations of leukocytes described by flow cytometry. Urine from 3 bladder cancer patients (P51-P53) treated with BCG was collected before the 6^th^ instillation (week 6 of treatment) and cells were analysed by flow cytometry in parallel. 3 populations could be identified in the FSC/SSC plots corresponding to CD15^+^ cells, CD14^+^ cells and CD14^+^CD15^+^ cells. However, different proportions of these populations were detected in each patient. These three populations were positive for CD16 although with different intensities and had different levels of expression for HLA-DR and CD11b.

To further confirm the lineage of urinary cells found by CyTOF and flow cytometry, gene expression was tested by qPCR. Dry cell pellets were used to extract RNA and, after cDNA synthesis, qPCR experiments were performed using primers for the main lineage markers, comparing their amplification cycle with that of the reporter gene, β-actin. To validate the primers, gene amplification from PBMC or purified granulocytes was performed and used for comparison (Figure 6). CD14 and CD66b expression were clearly detected in most of these samples, while CD3 and CD56 were only detected in a few patients and samples. It is interesting to note that CD66b was amplified more efficiently in cells from urine than in purified neutrophils from peripheral blood. This probably indicates that the former are active cells while granulocytes are usually quiescent and short lived in healthy donor peripheral blood. It is also possible to detect by PCR some markers that were not so evident by flow cytometry or CyTOF, suggesting the loss of epitopes or cell death.

**Figure 6.**
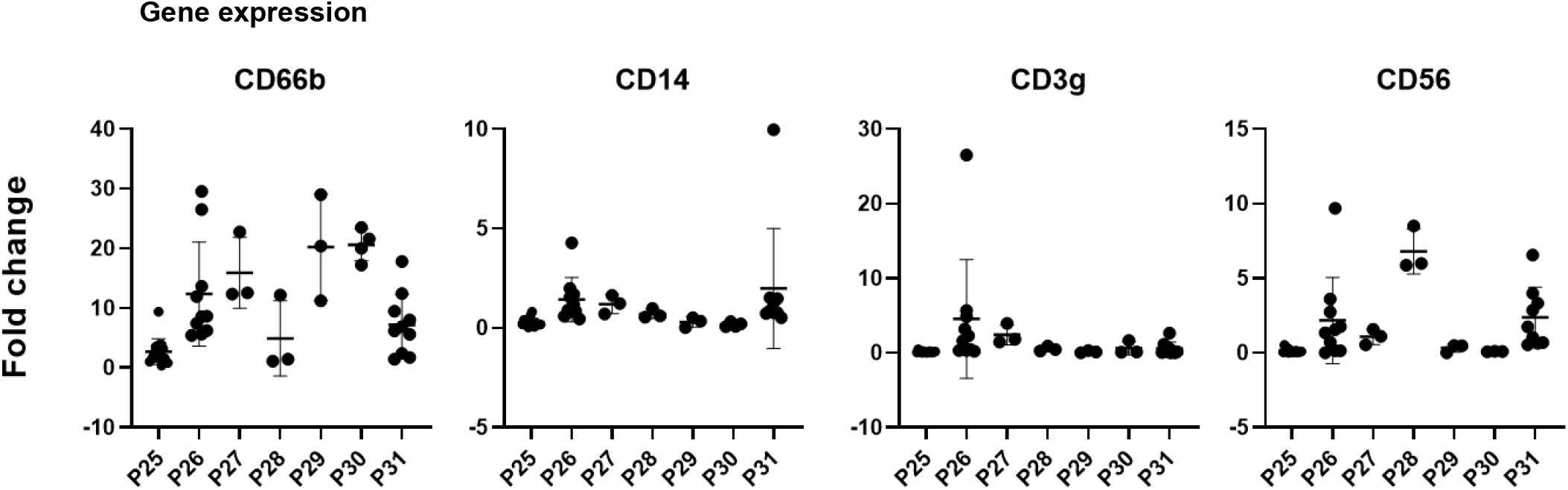
Gene expression analysis. Dry pellets from urine cells were processed for RNA extraction, followed by reverse transcription and cDNA were analysed by qPCR. Primers for the indicated genes were used and number of amplification cycles was measured. Plots represent the fold change relative to the reporter gene b-actin of a control population of granulocytes or PBMC (2^-ΔΔCt^).

### Urinary cell distribution can identify patients with unusual treatment outcomes

Notably, P25, P26 and P31 clearly had a different distribution to the rest of the patients. These three patients, in contrast with the majority of patients, already had immune cells in the sample collected at time zero, before starting BCG instillations (Figure 3) and had higher number of cells collected during the course of treatment (Supplementary Table 1). Interestingly, these three patients had different clinical course compared to the rest. P25 developed uveitis and died due to a brain haemorrhage. Whether this could be a case of a rare complication due to BCG is unclear; there are 15 cases of ocular inflammation reported after intravesical BCG instillations, 4 of them with confirmed Mycobacterial invasion (Liu *et al*, 2019). The treatment course of P26 was complicated by fever and urinary tract infection. P31 also had a gynaecologic tumour and thus had received radiotherapy. The composition of immune cells was also different in these three patients. P25 had a large NK cell population at time zero, that disappeared after BCG instillations, while P26 and P31 mainly had granulocytes at time zero. P26 mainly had high percentage of monocytes during the course of the treatment while P31 had large populations of both classical and APC-like granulocytes.

## DISCUSSION

Here, we report the first CyTOF analysis of cells released into urine in the context of BCG treatment for bladder cancer. We have also optimized flow cytometry protocols for the analysis of these cells and compared these data with qPCR analysis. With the aim of studying immune infiltration in bladder cancer patients, using a minimally invasive procedure, we analysed the cells in 44 urine samples obtained from 12 different bladder cancer patients at different time points during treatment. The majority of these samples were collected 3-7 days after receiving the previous BCG instillation, so that the initial inflammation response was already cleared. This study has allowed the characterization of several immune cell types that could be involved in the immune/inflammatory response in these cancer patients, including an APC-like neutrophil population.

Several technical challenges had to be overcome for this study, and data interpretation had to take into account several factors: 1) cells had been under hypoxic conditions and osmotic stress for several hours before collection, thus, certain epitopes might have been lost and autofluorescence increased; and 2) there is no healthy donor control, because urine only contains scarce cells in healthy donors, thus the methodology had to be set up comparing with PBMC. In addition to these problems, it was impossible to anticipate either the proportion of cell subpopulations expected or the epitopes that could be altered due to the cancer microenvironment.

The use of CyTOF allowed us to perform a broad screening of the cells released into urine because it is a highly multiplexed technique that permits the analysis of multiple markers on a limited number of cells, thus increasing the amount of information obtained in each experiment. Importantly, CyTOF is not affected by autofluorescence, a recurrent problem in cells that were obtained from urine and the technique provides many internal controls for non-specific binding, and so, we could demonstrate that the different antibodies used indeed bound to different cell populations in these samples.

This work provides a pilot comparative analysis of different samples (different time and different patients) revealing a striking cellular heterogeneity between urinary cells of BGC-treated patients. In general, activated immune cells with expression of CD107a and CD69 were identified and a shape consistent with activation (uropod and pseudopod formation) was also observed in microscopy. The myeloid lineage was also abundant, but different phenotypes were detected in different samples, revealing a high heterogeneity between patients.

A striking finding from these analyses is the presence in many samples of a population of cells expressing markers of both granulocytes and monocytes, reminiscent of other cell subtypes that have been defined in inflammatory or tumour microenvironments. One hypothesis was that CD14^+^CD15^+^ cells could be MDSC, which derive from a common hematopoietic progenitor cell and can express CD14 and CD15 (Bronte *et al*, 2016; Bruno *et al*, 2019; Gabrilovich, 2017). In particular, non-classical CD14^+^CD15^+^ MDSC have been described in lung and colon cancer (Mandruzzato *et al*, 2009; Vasquez-Dunddel *et al*, 2013). However, with this work, we used an optimized flow cytometry protocol, informed by our CyTOF screening, to demonstrate that CD14^+^CD15^+^ cells express markers of both neutrophils and APCs, likely representing a cell type described as antigen-presenting cells (APC)-like hybrid tumour associated neutrophils (TANs). APC-like hybrid TANs have been proposed to differentiate from classical neutrophils by exposure to tumour-derived cytokines, including IFN-γ and GM-CSF, and to possess enhanced antitumour properties, like antigen presentation and cross-presentation (Keerthivasan *et al*, 2014; Singhal *et al*, 2016). In the bladder cancer patient urine analysed here, CD15^+^CD14^+^ cells also expressed CD16, CD11c and HLA-DR. In flow cytometry CD14^+^ monocytes were easily distinguished from the double positive CD14^+^CD15^+^ granulocytes (putatively TAN). Whether this population could also have anti-tumour activities has not been addressed and should be the goal of future experiments. In these proposed new investigations, functional studies to distinguish between the activating or suppressor role of these cells will be performed. We propose that these populations could help to define new markers that aid the classification of patients as responder or non-responder, perhaps even to detect early the development of unwanted effects like infection spread. Thus, in new studies using the panels that we set up here, flow cytometric evaluation of the impact of these populations in disease outcome should be possible in a suitable cohort of patients.

Notably, lymphocytes were not abundant in these samples. Several factors could explain this: 1. Lymphocytes could still be attached to the bladder mucosa; 2. Lymphocytes could die in urine or not be recruited to urine; 3. Epitopes could be lost. Indeed, we have observed that certain clones of anti-CD3 antibodies lose the ability to recognise T cells after they are incubated in urine for 2 hours. Further, we could amplify CD56 and CD3 genes in some pellets of urine cells, suggesting that some subjects have lymphocytes present. CD3^+^ T cells found in urine samples were usually CD4^+^ although in some patients CD8 could also be detected. Further supporting the idea that some epitopes may be lost in urine, CD15 was not detected as easily as CD66b by flow cytometry even though both markers should stain the same population, and CD14 was not as bright in CyTOF as it was by flow cytometry. Together, these data suggest that antibody staining can be affected in urine samples.

Our analyses of qPCR confirmed the results obtained by CyTOF. We could identify three subjects who showed distinct cell population distribution compared to other patients. In general, patients responding well to the BCG therapy had fewer cells at the beginning of the treatment and developed primarily granulocyte populations in further weeks. This suggests that the composition of urinary cells may be related to the clinical outcome of the patient and could predict response to treatment; however, a larger study is needed in order to definitively establish the value of such biomarkers.

In this study, we have developed protocols that allow the follow-up of bladder cancer patients using flow cytometry of urine cells and gene expression analysis by PCR. Data complement our previous findings showing that cytokines are detectable 7 days after instillations with BCG, and that patients releasing very high amounts of CXCL10 had a more exacerbated immune response sometimes detrimental for the continuation of the treatment (Ashiru *et al*., 2019). We propose that new methods to evaluate the response to BCG bladder cancer patients will be possible using the analysis of cells and soluble factors in combination.

Further, flow cytometry and PCR are standard techniques used in every hospital for blood analysis that would make possible the study of high numbers of patients.

In conclusion, although we could not definitively establish links between treatment outcome and donor phenotypes in this challenging pilot study, we were able to identify atypical leukocyte populations possibly related to immune cells that have been found in other tumour microenvironments. This work establishes that it is possible to analyse cells released in patient urine by both flow cytometry and CyTOF and that these analyses can provide valuable information on both the intensity and the quality of the immune response leading to either clearance of the tumour or bacterial spread.

## MATERIALS AND METHODS

### Patient recruitment and study approval

A prospective study to analyse the immune response of BCG-treated NMIBC patients was designed and, a total of 44 urine samples were collected at different times of treatment from patients at either Hospital Infanta Sofía or Hospital Universitario La Paz (Madrid, Spain), n=14; mean age 72 years (Figure 7) (Table 1). Enrolment occurred between the years 2015 and 2016 and patients with bladder cancer at the stage Ta/T1 G3 or CIS, at diagnosis, were selected. Follow-up was done for an average of 2.2 years. No patient had recurrence. P26 suffered urine infection; P25 died due to brain haemorrhage; P46 died due to lung cancer; P43 and 48 had low number of cells in the only sample obtained from these patients, so they were not analysed. Because immune cells are not present in urine of healthy donors or mitomycin C-treated bladder cancer patients (our usual control), no control group was included. The experiments were conducted with the understanding and written informed consent of each participant and approved by local and regional ethical committees (CEI La Paz Hospital HULP-1067 Feb 8th 2011; revised by CEI Infanta Sofía Hospital and CSIC Local Ethical Committee; HULP-2338, April 25^th^ 2016). The experiments conformed to the principles set out in the WMA Declaration of Helsinki and the Department of Health and Human Services Belmont Report.

**Figure 7.**
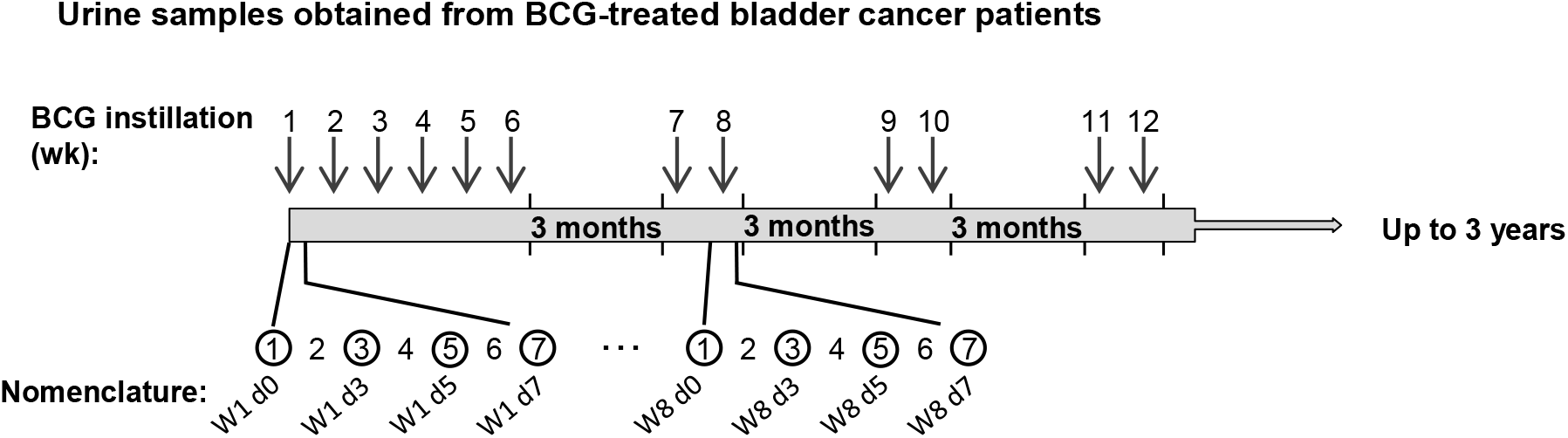
Schematic representation of the BCG treatment time line and urine samples obtained from bladder cancer patients. During the first cycle, patients received one weekly instillation for 6 weeks. After a three-month rest, they received a second cycle of two weekly instillations and this treatment continued up to three years. Arrows indicate the time (weeks) of the instillation calendar and circles an example of sample collection. The nomenclature used refers to the instillation week and the number of days elapsed since the instillation. For most patients, samples were obtained just before the instillations at week 1 (time 0, before starting the treatment), 3 and 6, as well as after the 3-month rest. In all these cases (except time 0), exposure to BCG occurred 7 days before. Samples were obtained 3 and 5 days after BCG for a few patients.

### Sample handling

Samples were obtained by spontaneous micturition by patients prior to BCG instillation at the clinic. Usually, within 2 hours, samples were centrifuged at 400 x g to separate cells and crystals from supernatants and washed with PBA [PBS containing 0.5% bovine serum albumin (BSA), 1% foetal bovine serum (FBS) and 0.1% sodium azide]. Cells were resuspended in PBA, counted and one aliquot was stained for flow cytometry immediately. The rest of the cells were washed by centrifugation at 200 x g and cryopreserved in FBS containing 10% DMSO at −80°C in aliquots until further analysis.

### Fluorescence microscopy

Cells were allowed to adhere on poly-L-lysine coated slides. Attached cells were then fixed with 4% p-formaldehyde for 10 min at RT, and stained with DAPI during 10 min at RT. After washing and mounting they were visualised using a DMI 6000B (Leica) photonic microscope at the confocal microscopy unit of the CNB.

### Electron microscopy

Washed urine cells were resuspended in 0.4 M HEPES buffer pH 7.2 and centrifuged at 200 x g before being fixed using 2% glutaraldehyde/1% tannic acid in HEPES buffer, 2 h at room temperature. The pellet was centrifuged to remove the fixing solution and resuspended in HEPES buffer and solidified by the addition of 2% agarose. After 1h incubation in 1% osmium tetroxide / 0.8% potassium iron cyanide at 4°C, and three washes, they were incubated with 2% uranyl acetate for 40 min at 4°C. After washing, samples were dehydrated using increasing percentages of acetone (30 % −100%), infiltrated in epon 812 and polymerised in BEEM™ embedding capsules for 48h at 60°C. 60-70 nm sections were obtained in a Leica EM UC6 ultra-microtome and supported in formvar/Carbon recovered Ni grids. Sections were stained with uranyl acetate and lead citrate and observed under a Jeol JEM 1011 a 100 kV microscope. Images were taken with a Gatan Erlangshen ES100W camera at the electron microscopy unit of the CNB.

### Flow cytometry

Cells washed in PBA were incubated with conjugated antibodies against surface markers at 4°C for 30 minutes in the dark [CD15 PB Biolegend #394704 Clone MMA, CD16 PC7 Biolegend #302016 Clone 3G8, CD14 A700 Biolegend #301822 Clone M5E2, HLA-DR PE Immunotech #1639 Clone IMMU-357, CD11c APC Biolegend #301614 Clone 3.9, CD3 APC Beckman coulter #IM2467 Clone UCHT1, CD19 ECD Beckman coulter #A07770 Clone J3-119, CD8 PC7 Biolegend #344712 Clone SK1, CD4 PE Immunotools #21278044 Clone EDU-2]. Cells were washed in PBA and analysed using either Gallios or CytoFLEX Flow Cytometers (Beckman Coulter). For urine flow cytometry, certain antibody clones did not work, so they were carefully selected; for example, CD3 OKT3 clone was changed for UCHT1. Analysis of the experiments was performed using Kaluza software (Beckman Coulter). In certain experiments, to keep cells intact after staining, they were fixed with 1% p-formaldehyde for 10 min at room temperature (RT).

### Mass Cytometry

Urine cells were thawed into 9 ml of 10% FCS-RPMI medium supplemented with 10 IU/ml Benzonase (Novagen 70664-3), counted and viability determined by trypan blue exclusion. Staining was performed in 96-well deep-well plates (Sigma) as previously reported (Strauss-Albee *et al*, 2015), briefly, 1-2·10^6^ cells were washed with 600 μl CyFACS (PBS supplemented with 2% BSA, 2 mM EDTA, and 0.1% sodium azide) and resuspended in 200 μL for staining cells were first incubated with 25 μM cisplatin (Enzo Life Sciences), then stained with a panel of surface antibodies (Supplementary Table 2). After that, cells were fixed and permeabilised (BD FACS Lyse, BD FACS Perm II), followed by intracellular antibody staining. Finally, cells were incubated with Ir-intercalator (DVS Sciences) and washed extensively using filtered PBS and metal free milli-Q water. Calibration Beads (EQ™ Four Element CB) were added before the sample was acquired in a mass cytometry instrument (Fluidigm). Antibodies were conjugated using MaxPar X8 labeling kits (DVS Sciences).

### Data normalization, viSNE analysis and hierarchical clustering

To ensure the homogeneity of data obtained at mass cytometry across time, data normalization was performed using Normaliser v0.3 [https://github.com/nolanlab/bead-normalization/wiki/Installing-the-Normalizer] (Finck *et al*, 2013).

Data were visualized with viSNE, implemented in Cytobank (https://www.cytobank.org/) (Amir el *et al*., 2013).

Several viSNE runs were performed grouping different samples (either according to patient or time of treatment). Selection of populations within viSNE was done as follows: live cell populations were first selected gating intact cells based on 191Ir (DNA-1) vs 193Ir (DNA-2), then live singlets were gated based on the parameters corresponding to event length, beads and cisplatin (Figure 2A) [26]. For certain analyses, indicated in the main text, a further gating was performed to select CD66b^+^ cells. Equal numbers of cells (100,000) were selected.

After gating, viSNE analyses were launched selecting the channels for the population separation. Most analyses were performed selecting the lineage markers: CD19, CD4, CD8, CD3, CD15, CD14, NKp46, CD66b, CD56, HLA-DR and CD33. In the case of the analysis performed after gating the CD66b^+^ population, the parameters selected to launch the viSNE analysis were CD11b, CD35, CD63, CD11c, CD14, CD107a. Different populations were manually gated on t-SNE1 vs t-SNE2 plots and median expression of the different markers was used to group clusters with similar expression profiles.

Hierarchical cluster of arrays was performed using cluster 3.0, selecting the average linkage clustering method. Java TreeView was used for visualization, heat map was generated in GraphPad.

### qPCR

Urine was centrifuged and the cell-containing pellet was washed with PBS. RNA was extracted from dry pellets using RNeasy Kit (Qiagen). cDNA synthesis was carried out using random hexamers and SuperScript II RNase (Invitrogen). For qPCR, 3 μl of cDNA (diluted 1:10) were mixed with 5 μl of reaction mix [PyroTaq EvaGreen qPCR Mix Plus (ROX) (CMBTM) and both forward and reverse primers at 0.5 mM] in 384-well plates. Wells containing 3 μl of water instead of cDNA were used as negative controls. For each amplification, b-actin was included as reporter gene. Amplification reactions were set in triplicates and performed in a QuantStudio5 (Applied Biosystems) instrument with a first cycle of 10 min at 95°C followed by 40 cycles of amplification (15 s at 95°C and 1 min at 60°C). Results were analysed with QuantStudio™ Design & Analysis 1.4.3 Software and Microsoft Excel. Fold change between a control population (purified neutrophils, for CD66b; PBMC, for the other genes) and the urine sample were calculated relative to the cycles of b-actin (2^-ΔΔCt^). Plots were represented using GraphPad Prims 8 Software (GraphPad Software, USA, www.graphpad.com). The following primers were used: CD56 Forward: TCTGGATGGGCACATGGTG; CD56 Reverse: TGCTCTTCAGGGTCAGCGA; CD3g Forward: GGGATGTATCAGTGTAAAGG; CD3g Reverse: CAGCAATGAAGTAGACCC; CD14 Forward: ACGCCAGAACCTTGTGAGC; CD14 Reverse: GCATGGATCTCCACCTCTACTG; b-actin Forward: CCCAGCACAATGAAGATCAA; b-actin Reverse: CGATCCACACGGAGTACTTG; CD66b Forward: GCGAGTGCAAACTTCAGTGACC; CD66b Reverse: ACTGTGAGGGTGGATTAGAGGC.

## Author contributions

EC, CS, LS, EV, TR, HTR and MVG performed experiments and analysed data. GE, EMGC, SLC, MAM, ALi, ALe and MVG selected, acquired and processed patient samples. LMP, CB and MVG provided material support and supervised the study. CB, MVG and LMP conceived and designed the study; EC, LMP, CB and MVG wrote the manuscript.

## The Paper Explained

### Problem

BCG treatment of NMIBC patients is a successful example of immunotherapy, however, there are still many questions on how the inflammation process results in tumour elimination. To date, no systematic longitudinal information is available on the cells recruited to the bladder of BCG-treated patients in samples obtained after the initial cytokine cascade has finished. Since the identity of the cells appearing in urine several days after each instillation can provide information on the intensity and the quality of the immune response, a prospective study was planned to establish whether it would be possible to characterize immune cells by cytometry methods and to establish baseline values to facilitate follow-up studies.

### Results

Different patients had different numbers of cells in urine and diverse percentages of immune populations could be identified. As previously reported, granulocytes were very abundant, however, detailed phenotypic analysis revealed that these cells might mediate either activation or suppression in different patients. A particular phenotype, reminiscent of tumour associated neutrophils, easily detectable by flow cytometry has been identified.

### Impact

This first detailed analysis of cells in urine, using a sophisticated laboratory-based methodology, has also been optimised for translation to standard techniques employed in every hospital for blood analysis. Thus, it provides the proof-of-concept for the follow-up of bladder cancer patients using urine cells and for early identification of patients with an excessive, detrimentally strong or insufficient response to BCG therapy.

## Acknowledgements

The authors would like to thank the collaboration of patients and nurses from Infanta Sofía and La Paz Hospitals; the confocal microscopy, electron microscopy and flow cytometry services at the CNB; all the members of the Blish’s laboratory for help and advice on CyTOF experiments.

This work was supported by grants from Madrid Regional Government “IMMUNOTHERCAN” [S2017/BMD-3733-2 (LMP, MVG)]; the Spanish Ministry of Science and Innovation [RTI2018-093569-B-I00, RTC-2017-6379-1 (MCIU/AEI/FEDER, EU) (MVG)]; CS, EMGC and SLC were recipients of Fellowships from Erasmus+, Fundación La Caixa and Spanish Ministry of Education (FPU) respectively. MVG was a Visiting Associate Professor at the University of Stanford funded by a “Salvador de Madariaga” grant, Spanish Ministry of Education (MECD) and NIH DP2AI112193 (CAB).

**Supplementary information is available at ‘EMBO Molecular Medicine’s website’**

---

**Supplementary Table 1.** Cell count in urine.

| Patient | Week | Days after instillation | x10<sup>3</sup> cells/ml |
| --- | --- | --- | --- |
| P25 | Before first instillation |  | 9 |
|  | w1 | d2 | 64 |
|  |  | d5 | 90 |
|  |  | d7 | 33 |
|  | w2 | d7 | 95 |
|  | w3 | d7 | 75 |
|  | w5 | d7 | 1200 |
|  | w6 | d5 | 720 |
|  |  | d7 | 2619 |
|  | w19 | d2 | 75 |
|  |  | d5 | 53 |
|  |  | d7 | 22 |
| P26 | Before first instillation |  | 40 |
|  | w1 | d5 | 159 |
|  |  | d7 | 94 |
|  | w2 | d7 | 80 |
|  | w3 | d2 | 100 |
|  |  | d7 | 30 |
|  | w6 | d5 | 18 |
|  |  | d7 | 9 |
|  | w7 | d2 | 64 |
|  | w18 | d7 | 4 |
| P27 | Before first instillation |  | 0 |
|  | w2 | d7 | 10 |
|  | w5 | d7 | 43 |
| P28 | Before first instillation |  | 0 |
|  | w2 | d7 | 60 |
|  | w5 | d7 | 188 |
| P29 | Before first instillation |  | 0 |
|  | w2 | d7 | 9 |
| P30 | Before first instillation |  | 0 |
|  | w2 | d7 | 10 |
|  | w3 | d2 | 248 |
|  |  | d7 | 94 |
|  | w6 | d2 | 200 |
|  |  | d7 | 9 |
| P31 | Before first instillation |  | 4800 |
|  | w1 | d2 | 1660 |
|  |  | d7 | 1510 |
|  | w2 | d7 | 720 |
|  | w3 | d2 | 1089 |
|  |  | d7 | 750 |
| P41 | w5 | d7 | 4 |
| P42 | w2 | d7 | 12 |
| P44 | w5 | d7 | 12 |
| P45 | w5 | d7 | 12 |
| P46 | w2 | d7 | 12 |
| P47 | w5 | d7 | 60 |

**Supplementary Table 2.** CyTOF antibody panel.

| Isotope | Ab | Clone | Company |
| --- | --- | --- | --- |
| Qdot - Cd | HLA-DR | Tu36 | Life technologies |
| 89Y | EpCAM | 9C4 | Biolegend |
| 115In | CD19 | HIB19 | Biolegend |
| 141Pr | CD38 | HIT2 | BioLegend |
| 142Nd | CD11b | ICRF44 | Biolegend |
| 143Nd | CD4 | OKT4 | Biolegend |
| 144Nd | CD8 | SK1 | Biolegend |
| 145Nd | CD57 | HCD57 | Biolegend |
| 146Nd | PD-1 | EH12.2H7 | Biolegend |
| 147Sm | CD3 | UCHT1 | Biolegend |
| 148Nd | CD35 | E11 | Biolegend |
| 149Sm | TLR2 | TL2.1 | Biolegend |
| 150Nd | CD45 | HI30 | Biolegend |
| 151Eu | CD15 | HI98 | Biolegend |
| 152Sm | CD63 | H5C6 | Biolegend |
| 153Eu | CD11c | Bu15 | Biolegend |
| 154Sm | CD14 | M5E2 | Biolegend |
| 155Gd | NKp46 | 9E2 | Biolegend |
| 156Gd | TCRgd | B1 | Biolegend |
| 157Gd | NKG2C | 134522 | R&D System |
| 158Gd | CD24 | ML5 | Biolegend |
| 159Tb | CD7 | CD7-6B7 | Biolegend |
| 160Gd | CD69 | FN50 | Biolegend |
| 161Dy | CD66b | G10F5 | Biolegend |
| 162Dy | CD64 (FcgrI) | 10,1 | Biolegend |
| 163Dy | Invariant NKT | 6B11 | Biolegend |
| 164Dy | CD89 | A59 | Biolegend |
| 165Ho | CD107a | H4A3 | Biolegend |
| 166Er | TLR4 | HTA125 | Biolegend |
| 167Er | CD94 | DX22 | Biolegend |
| 168Er | MICA | MAB13001 | R&D System |
| 169Tm | CD32 (FcgrIIa) | FUN-2 | Biolegend |
| 170Yb | CD107b | H4B4 | Biolegend |
| 171Yb | NKG2A | 131411 | R&D systems |
| 172Yb | CD25 | M-A251 | Biolegend |
| 173Yb | CD95 | DX2 | Biolegend |
| 174Yb | CD56 | NCAM16.2 | BD |
| 175Lu | CD33 | WM53 | Biolegend |
| 176Yb | HLA-DQ | Tu169 | Biolegend |
| 209Bi | CD16 (FcgrIII) | 3G8 | Fluidigm |

**Supplementary Figure 1.**
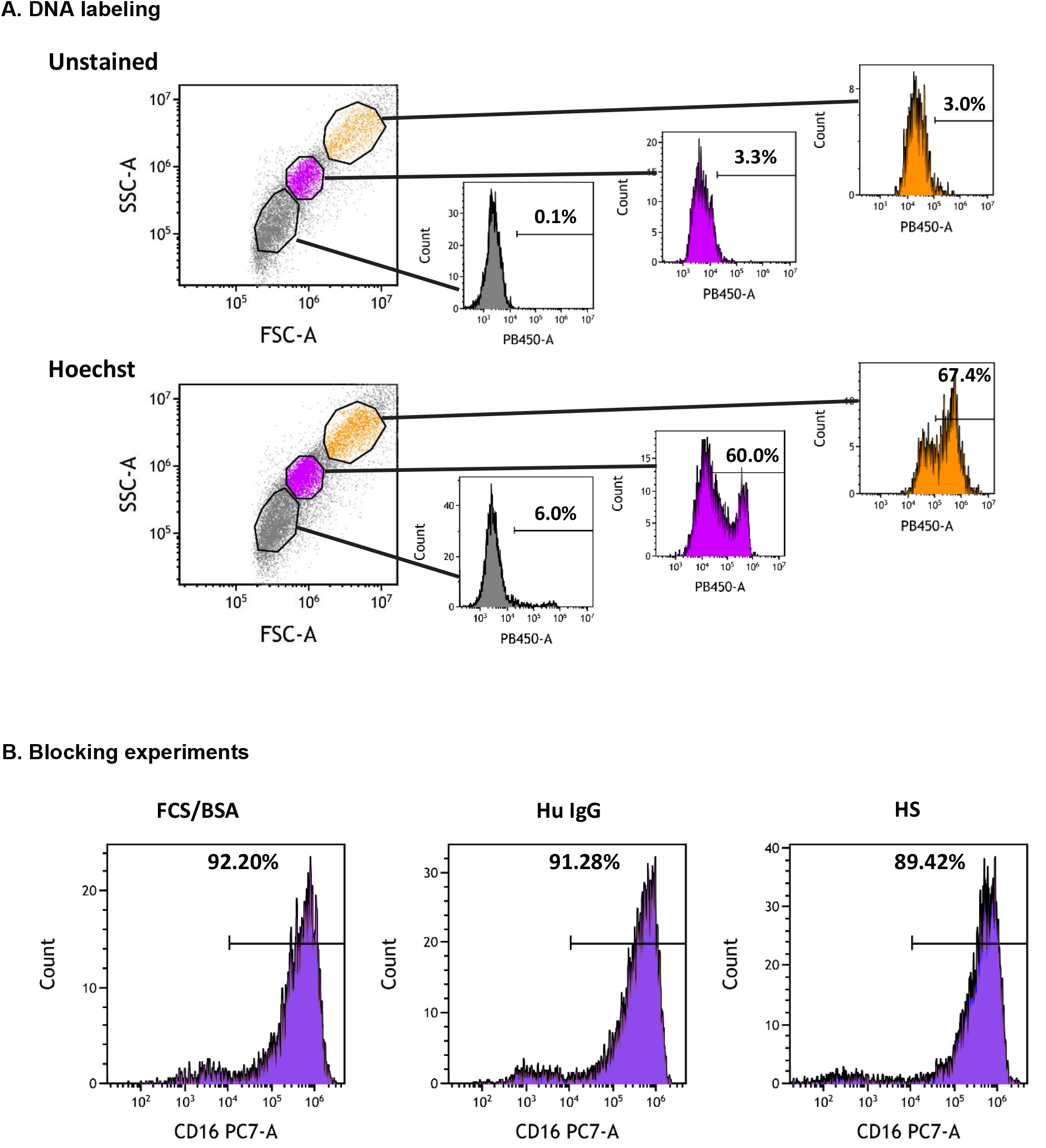
Flow cytometry analysis of urine cells. **A. Flow cytometry gating strategy and DNA labeling**. Doublets were excluded in the FSC_height_/FSC_area_ plots. Hoechst staining was analysed within three different gates in FCS/SSC as depicted. Upper panels correspond to unstained samples and lower panels to samples stained with 1/100 Hoechst 33342 dye. The percentage of Hoechst positive cells is shown for each histogram; the position of the marker was maintained in each region. The grey region corresponds to DNA-negative events. **B. Blocking experiments**. Different conditions for blocking of Fc receptors were tried: PBA [PBS supplemented with 0.5% bovine serum albumin (BSA), 1% FBS and 0.1% sodium azide], 10 mg/ml of purified human IgG and 10% human serum (HS). Cells were stained with antibodies against CD14 and CD16. The figure represents histograms of the CD16 staining within the CD14 positive cells. All the blocking conditions have similar staining pattern.

**Supplementary Figure 2.**
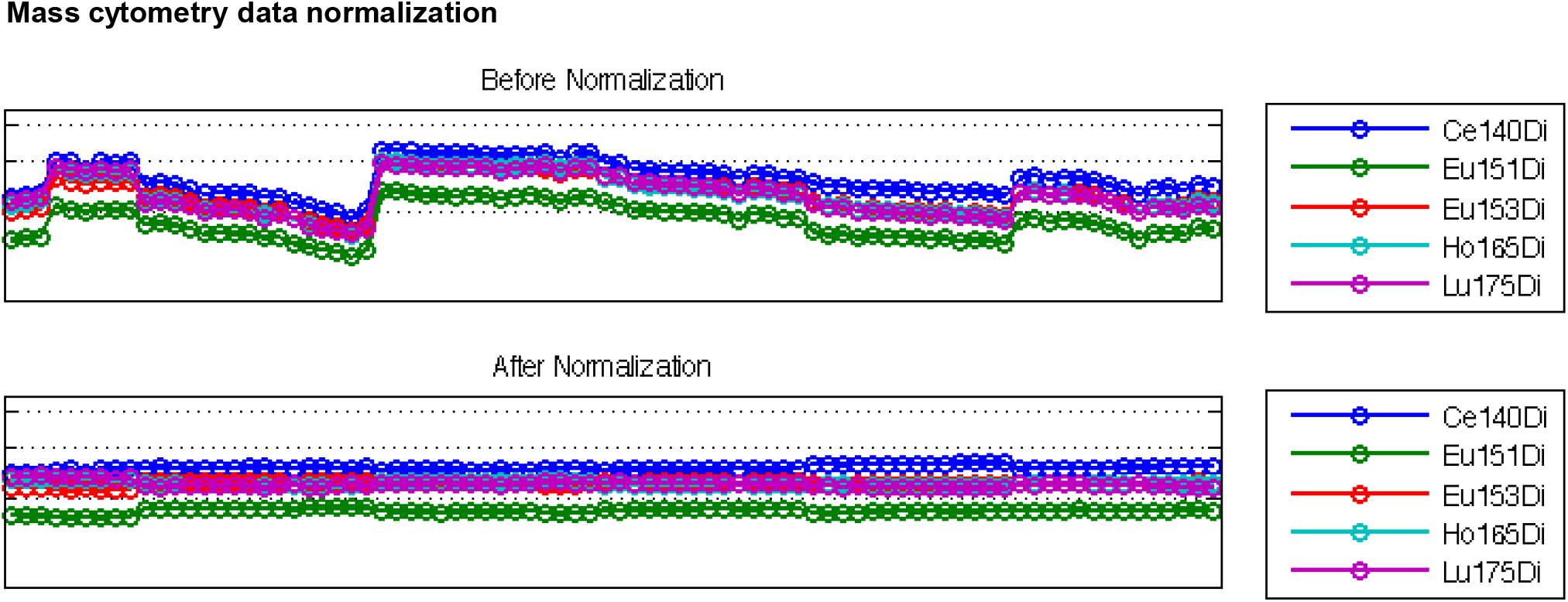
Mass cytometry data normalization. Data collected in all CyTOF experiments were run through Normaliser v0.3.

**Supplementary Figure 3.**
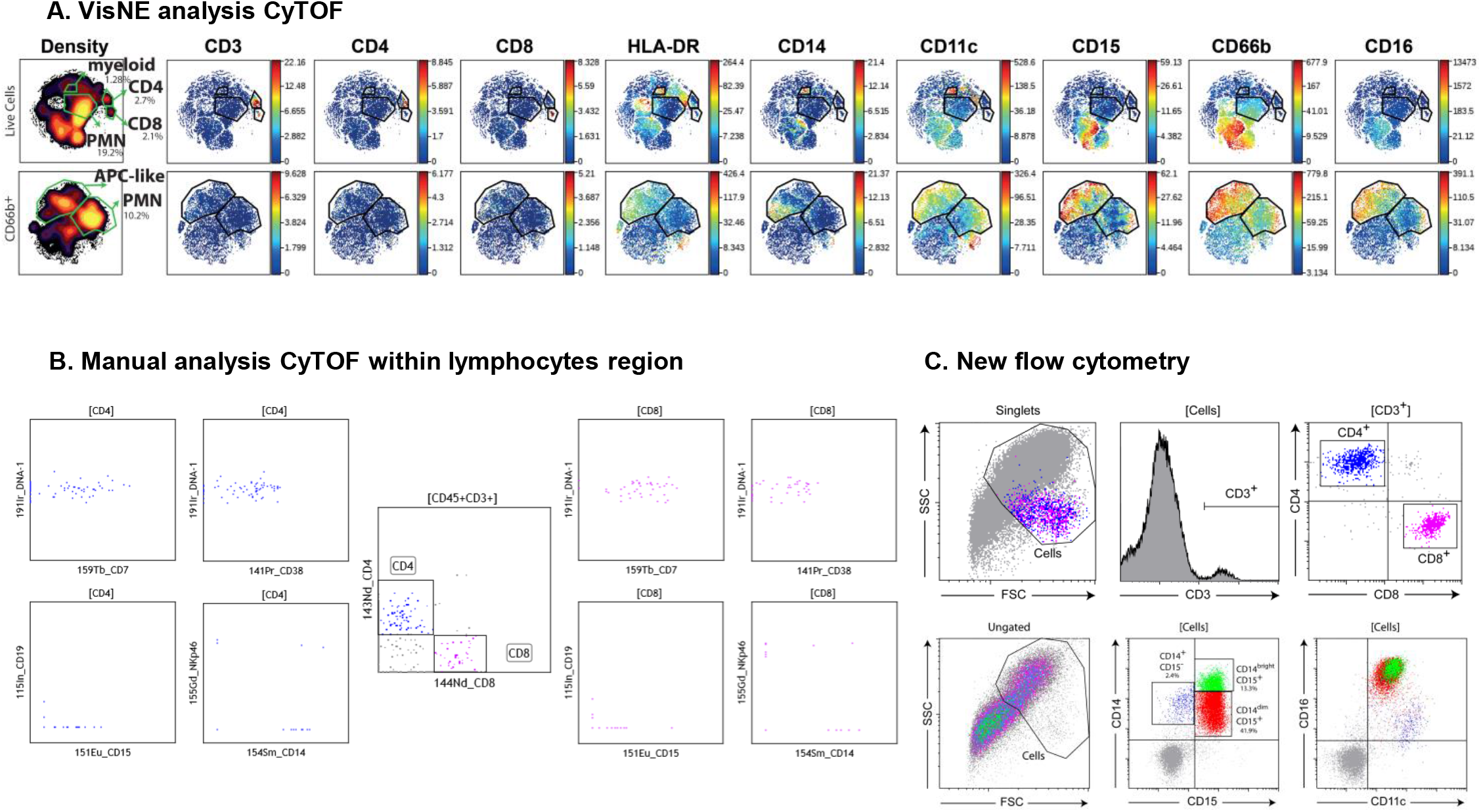
Leukocytes released to urine analysed by CyTOF and flow cytometry. **A. viSNE analysis.** CD3^+^ cells could be visualised in viSNE analyses and these lymphocytes could be classified as CD4 or CD8. Similarly, CD66b cells, also positive for CD15, were identified. **B. Manual analysis of CyTOF data.** CyTOF data were analysed using flow cytometry software to identify CD3 positive cells, CD4^+^ and CD8^+^. **C. Flow cytometry.** Cells from the same patient were analysed by flow cytometry to confirm the presence of CD3 lymphocytes. A population of CD15^+^CD14^+^CD16^+^ cells was confirmed.

**Supplementary Figure 4.**
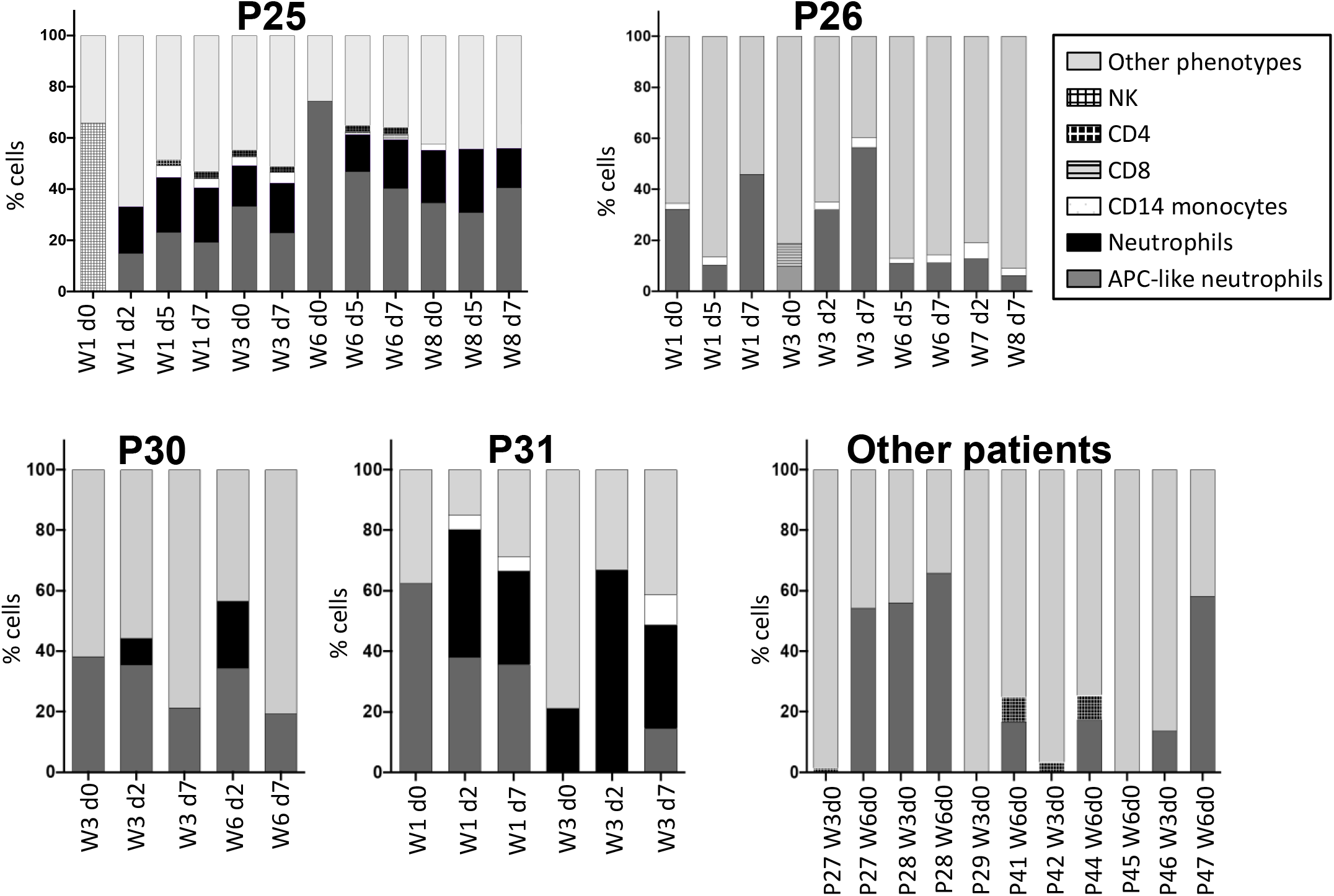
Immune populations in different patients. Graphs represent the number of cell populations found in each patient in viSNE analysis of data obtained in CyTOF.

## Notes

**Conflict of interest** The authors declare no potential conflict of interest.

